# Epstein-Barr virus promotes survival through germinal center light zone chromatin architecture

**DOI:** 10.1101/2020.10.22.350108

**Authors:** Joanne Dai, Emma Heckenberg, Lingyun Song, Gregory E. Crawford, Micah A. Luftig

## Abstract

BFL-1 is an understudied anti-apoptotic protein upregulated in cancer and Epstein-Barr virus (EBV)-immortalized lymphoblastoid cell lines (LCLs). We have previously shown that BFL-1 is regulated through viral EBNA3A-mediated alterations in B-cell chromatin conformation (Price et al., 2017). Here, we extend those findings to define cis- and trans-acting factors that regulate BFL-1 in LCLs and reliance on BFL-1 for survival from extrinsic apoptosis. Beyond LCLs, BFL-1 is expressed in B cells maturing in the germinal center (GC). We therefore characterized the gene expression profiles and chromatin landscape of maturing human tonsillar B-cell subsets. While chromatin accessibility at the BFL-1 locus increases as naïve B cells enter the GC reaction, BFL-1 expression increases during the transition from dark zone to light zone (LZ) correlating with association between enhancer regions and the transcriptional start site. The relationship between LCLs and LZ B cells suggests that EBV phenocopies GC biology to enhance their survival in establishing latent infection.

## INTRODUCTION

The Epstein-Barr virus (EBV) is a ubiquitous gamma-herpesvirus that infects >95% of the global adult population. EBV latent infection is life-long and is established in quiescent memory B cells, presumably by undergoing the germinal center reaction alongside maturing, uninfected B cells (Thorley-Lawson et al., 2013). In this model of infection, known as the Germinal Center Model, primary EBV infection is asymptomatic and escapes immune detection by robust T-cell mediated responses. However, in the absence of an immune response, such as in HIV infection or organ transplant, unchecked EBV infection can give rise to cancers of both B cell and epithelial cell origin. Worldwide, nearly 200,000 patients each year are diagnosed with EBV-positive cancer (Shannon-Lowe and Rickinson, 2019).

EBV-driven oncogenesis is modeled through the *in vitro* infection of human B cells, which leads to their growth transformation into immortalized lymphoblastoid cell lines (LCLs). This process requires the expression of six latent proteins: EBV nuclear antigen 1 (EBNA1), EBNA2, EBNA3A, EBNA3C, EBNA-LP and latent membrane protein1 (LMP1). The EBNAs are expressed early after infection and regulate viral and host gene expression by hijacking host transcriptional machinery and transcription factors. EBNA2 induces c-Myc expression that drives rapid hyperproliferation and induction of the DNA damage response that limits EBV-mediated transformation (Nikitin et al., 2010). Infected B cells in the early-phase, or approximately a week after initial infection, express the Latency IIb program, which is characterized by low LMP-1 expression. About 3-5 weeks post-infection, infected B cells have grown out into LCLs and express the Latency III program, which is characterized by full expression of all viral products (Price and Luftig, 2015; Price et al., 2012). This includes high levels of LMP-1, which mimics an active CD40 receptor that signals constitutively through NFκB and is required for proliferation and survival (Cahir-McFarland et al., 2000; Gires et al., 1997).

Our lab has found that the early- and late-phases of EBV are transcriptionally distinct (Messinger et al., 2019) and utilize unique strategies to promote apoptosis resistance (Price et al., 2017). Apoptosis is regulated by complex protein-protein interactions between pro-apoptotic and anti-apoptotic members of the BCL2 protein family. While uninfected B cells are dependent upon the anti-apoptotic protein BCL-2 for survival, early-infected B cells depend upon BCL-2 and MCL-1, and late-infected B cells at the LCL stage upregulate dependency upon BFL-1. We determined that upregulation of BFL-1, which is expressed from the *BCL2A1* gene, requires EBNA3A, a viral transcriptional cofactor that activates and represses viral and host gene expression by regulating enhancer activity and chromatin architecture (Bazot et al., 2015; Harth-Hertle et al., 2013; Hertle et al., 2009; McClellan et al., 2013). To promote BFL-1 transcription, EBNA3A promotes the looping of upstream enhancer regions to the transcriptional start site (TSS). In EBNA3A-null LCLs, these enhancer-promoter interactions are lost and levels of active transcription machinery at the TSS are reduced. In addition to its role in inhibiting expression of the pro-apoptotic protein BIM, EBNA3A promotes survival in EBV-immortalized LCLs through chromatin-mediated regulation of apoptotic proteins.

Like LCLs, germinal center light zone (GC LZ) B cells express high levels of BFL-1 mRNA induced by strong activation of the CD40 receptor (Victora et al., 2012); however, the non-viral mechanisms underlying BFL-1 upregulation remain unknown. BFL-1 expression is so high that it is frequently used as a marker to distinguish LZ GC B cells from other B lymphocytes (Milpied et al., 2018). Historically, the physiological role of BFL-1 was difficult to study because of gene quadruplication in mice, whereas humans express one gene (Hatakeyama et al., 1998). However, *in vivo* studies made possible by transgenic RNAi mouse models suggest that BFL-1 is important for the survival of activated, mature B cells (Ottina et al., 2012; Sochalska et al., 2015). This has not yet been testable in human GC B cells, which are especially sensitive to spontaneous apoptosis *ex vivo* and are genetically intractable, making it challenging to study BFL-1 upregulation in GC LZ B cells.

In this study, we follow up on Price *et al.*, 2017 and further characterize the chromatin architecture and enhancer-mediated BFL-1 transcription in EBV-immortalized LCLs. We find that this architecture strongly resembles that in GC LZ B cells, suggesting that EBV infection *in vitro* innately recapitulates certain aspects of the GC reaction. Because human GC LZ B cells are not amenable to genetic experiments, we use LCLs to perform functional analysis of BFL-1 enhancer-promoter interactions. Our analysis also reveals a large overlap of dynamically expressed gene targets in GC B cells and EBV-infected B cells. By characterizing the chromatin landscape of LCLs and GC LZ B cells, we present data that supports the long-standing Germinal Center Model of EBV infection *in vivo* that postulates that virus-infected B cells undergo the germinal center reaction to establish life-long latent infection.

## RESULTS

### Enhancer activity and chromatin-bound levels of YY1 are associated with BFL-1 transcription

Previously, we found that the levels of active transcription machinery and chromatin looping to the BFL-1 TSS were dependent upon EBNA3A binding to distal enhancers. To further characterize the chromatin landscape, we used publicly available ChIP-seq data performed on the GM12878 LCL as well as annotations predicted by ChromHMM (**Figure 1A**). ChromHMM is an automated computational system for learning, characterizing, and visualizing genome-wide maps of annotated chromatin states (Ernst and Kellis, 2012). Publicly available Chromatin Interaction Analysis by Paired-End Tag Sequencing (ChIA-PET) RNA Pol II data, which describes the long-range interactions of chromatin sites associated with transcription (Tang et al., 2015), also showed a high level of enhancer-promoter interactions between the BFL-1 TSS and upstream regions. We therefore decided to focus on the following regions: 1) Enhancer 1 (Enh 1), a nearby enhancer bound by NFκB transcription factors, EBNA2 and EBNA-LP, and elevated levels of H3K27ac and H3K4me1; 2) a RelA/B-binding site (RBS); 3) Enhancer 2 (Enh 2), a distal enhancer bound by RelA/B, H3K27ac and H3K4me1, and viral proteins EBNA-LP and EBNA3A; and 4) the EBV regulatory element (ERE), a putative EBV super-enhancer bound by all viral nuclear proteins.

**Figure 1.**
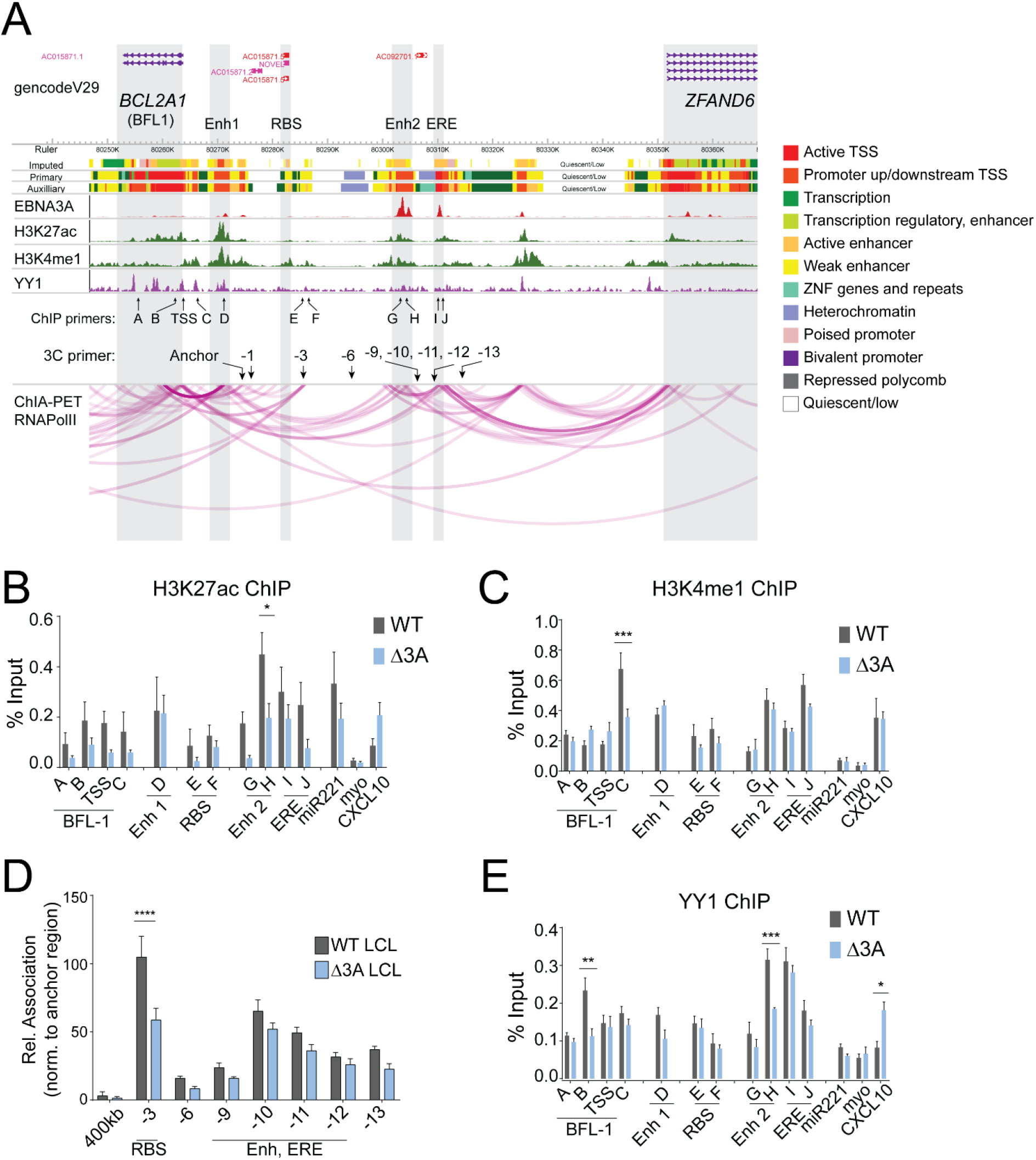
Increased enhancer activity and interactions to the BFL-1 TSS promotes BFL-1 transcription in WT LCLs. **A)** WashU Epigenome Browser screenshot of publicly available ChIP-seq and RNA Pol II ChIA-PET datasets performed on the GM12878 LCL, aligned to hg19 assembly, with labeled regions of interest and locations of primers used for ChIP-qPCR. ChromHMM annotations and tracks are included to show demarcated transcribed and active enhancer regions (Jiang et al., 2017). Relative positions of primers used for 3C-qPCR and ChIP-qPCR are included. **B)** H327ac and **C)** H3K4me1 ChIP-qPCR performed on WT and Δ3A LCLs. Regions corresponding to miR221 (EBNA3A-upregulated target), myoglobin (myo, not expressed in LCLs; negative control), and CXCL10 (EBNA3A-downregulated target) were also included. Enrichment calculated relative to input. Mean and SEM from three experiments are plotted. Significance determined by Two-way ANOVA. * p < 0.05, ** p < 0.01, *** p < 0.001. **D)** 3C-qPCR of interactions of HindIII fragments produced from WT and Δ3A EBV-immortalized LCLs. Interaction frequencies were normalized to those of the nearest neighbor HindIII fragment. Mean and SEM from three experiments are plotted. Significance determined by Two-way ANOVA with Holm-Sidak’s multiple comparisons test. *** p < 0.001. **E)** YY1 ChIP-qPCR performed on WT and Δ3A LCLs. Enrichment calculated relative to input. Mean and SEM from three experiments are plotted. Significance determined by Two-way ANOVA. * p < 0.05, ** p < 0.01, *** p < 0.001.

To determine the enhancer activities of these regions of interest, we performed ChIP-qPCR for H3K27ac and H3K4me1(Rada-Iglesias et al., 2011) (**Figure 1B-C**). Overall, H3K27ac levels were elevated in WT LCLs compared to ΔEBNA3A LCLs, especially at Enh 2. H3K4me1 levels were mostly similar between WT and ΔEBNA3A LCLs, although there was an increase in H3K4me1 near the BFL-1 TSS in WT LCLs that confirms our previous findings (Price et al., 2017). Reduced H3K27ac, but comparable levels of H3K4me1, at Enh 2 suggests that while this enhancer is adequately primed by bound chromatin regulators, its activity is dependent upon EBNA3A. Similarly, H3K27ac levels at the ERE did not change significantly in the absence of EBNA3A, suggesting that enhancer activity from this region is sufficiently maintained by the other EBNAs.

We next sought to understand the mechanism by which EBNA3A loss led to reduced chromatin interactions between the BFL-1 TSS and upstream enhancers (**Figure 1D**). Enhancers, both viral and non-viral, are characterized by heightened levels of chromatin-bound structural factors, such as YY1. YY1 is a transcription factor that facilitates enhancer-promoter interactions by dimerizing on chromatin (Weintraub et al., 2017; Zhou et al., 2015). Publicly available ChIP-seq data from LCLs indicate that YY1 binds at *BCL2A1* and associated regions of interest, suggesting that the chromatin architecture in WT LCLs is mediated by YY1 (**Figure 1E**). Indeed, when we performed ChIP-qPCR for YY1 in EBNA3A-null LCLs, we found that YY1 levels were reduced both distally at Enh 2 and within the BFL-1 gene body. This indicated that EBNA3A may play a role in recruiting YY1 in mediating long-distance chromatin interactions for BFL-1 transcription.

### BFL-1 transcription in LCLs results from combined activities of upstream genomic regions and enhancers

To determine if upstream regions were important for BFL-1 transcription, short-guide RNAs (sgRNAs) were targeted to the BFL-1 TSS, Enh 1, RBS, Enh 2, and ERE. These sgRNAs were co-expressed with dCas9-KRAB to repress transcription and enhancer activity (Thakore et al., 2015). Because RNA Pol II ChIA-PET in LCLs showed high levels of interconnectivity between these regions (**Figure 1A**), we surmised that constitutive targeted inhibition of an enhancer could strengthen or induce looping to nearby enhancers to rescue BFL-1 expression (Klein et al., 2018). We therefore utilized a Tetracycline (Tet; TRE)-inducible dCas9-KRAB system to repress targeted enhancers (Fulco et al., 2016; Kearns et al., 2014) (**Figure 2A**), such that doxycycline (Dox) treatment induced expression of dCas9-KRAB.

**Figure 2.**
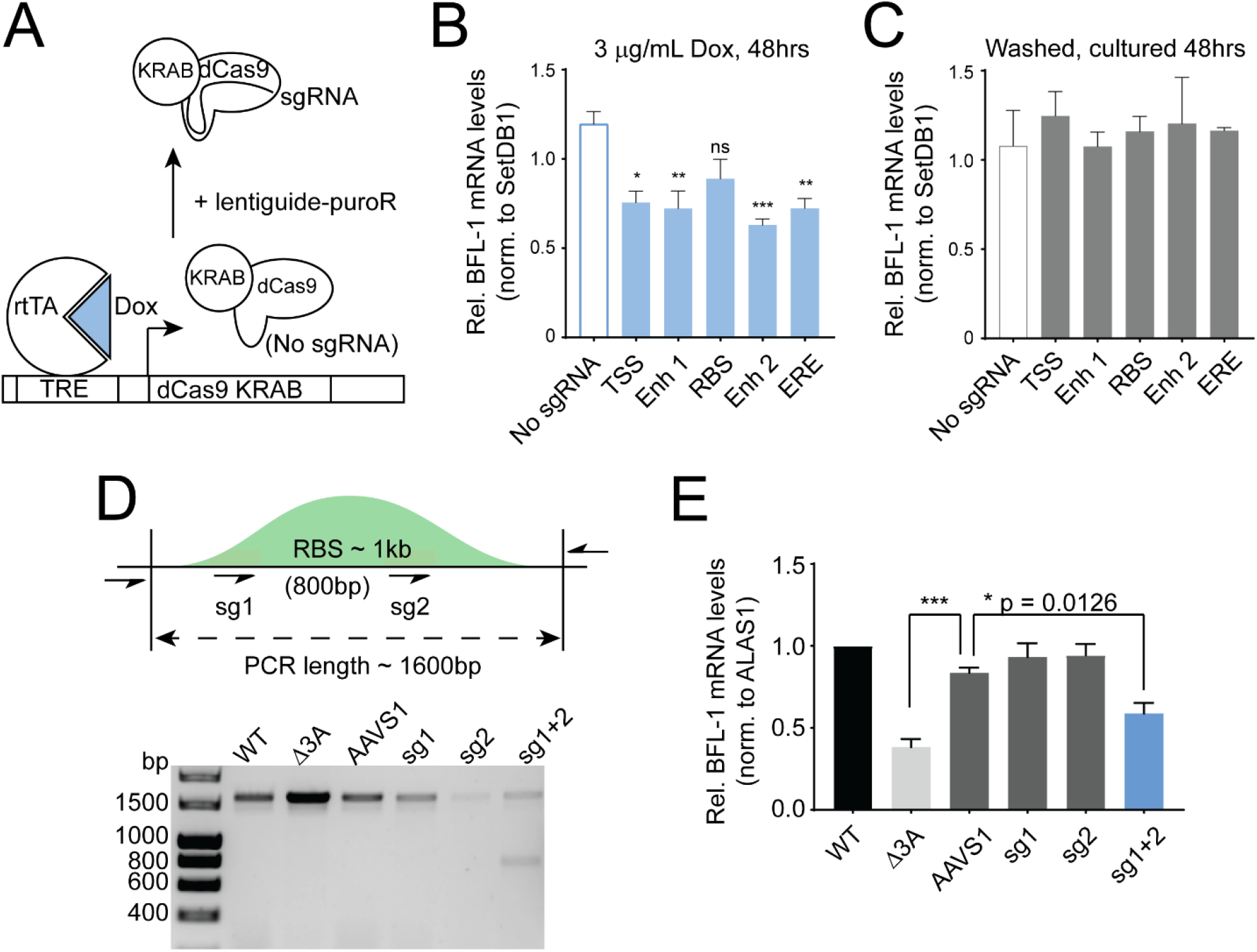
Upstream genomic regions are important for BFL-1 transcription. **A)** Schematic depicting TRE-dCas9-KRAB experimental system. Doxycycline (Dox) addition induces the expression of dCas9-KRAB but requires sgRNA expression from the lentiguide-puroR system to target the genomic DNA. **B)** qPCR performed on LCLs transduced with TRE-dCas9-KRAB with or without sgRNAs targeting the BFL-1 TSS and upstream regions. Cells were treated with 3mg/mL doxycycline (Dox) for 48 hours; in **C)** Dox-treated cells are washed and then treated for 48hrs. Mean and SEM are plotted and normalized to Untreated samples, significance determined through One-way ANOVA with multiple comparisons to “No sgRNA” control. **D)** DNA gel showing representative PCR confirming Cas9-mediated editing of the RBS region. PCR was performed on genomic DNA isolated from donor-matched WT and Δ3A LCLs and wildtype LCLs 5 days post-transfection with Cas9 coupled with sgRNAs targeting the AAVS1 site, sg1 and sg2 of the RBS, and combined sg1 and sg2. **E)** qPCR performed on WT and Δ3A LCLs and WT LCLs transfected with Cas9 RNPs targeting the RBS. Results are from 4 separate transfections on two separate days, day 5 post transfection. Significance by One-way ANOVA, multiple comparisons, mean and SEM reported. *p < 0.05, ***p < 0.005.

Dox-induced repression of the TSS, Enh1, RBS, Enh 2, and ERE led to reduced BFL-1 mRNA levels compared to a non-targeting control (**Figure 2B**), indicating that multiple regions contribute to BFL-1 expression in LCLs. Washing out doxycycline rescued BFL-1 knockdown, confirming drug-specific effects (**Figure 2C**). The TSS, Enh 1, Enh 2, and ERE are characterized by high H3K27ac peaks and were consequently more sensitive to repression by dCas9-KRAB, which causes histone methylation and deacetylation (Thakore et al., 2015).

Of the regions targeted, the RBS had relatively low H3K27ac levels, so while dCas9-KRAB targeted at the RBS led to an observable decrease in BFL-1 mRNA levels, this did not achieve significance. Nonetheless, because interactions between the RBS and the BFL-1 TSS fragments were so significantly enriched in WT LCLs, we hypothesized that the RBS was indeed important for BFL-1 transcription. We then used CRISPR/Cas9 to delete the RBS by using two sgRNAs that flanked the RelA and RelB ChIP-seq peaks (**Figure 2D**). Using both guides together generated a ~800bp deletion in the RBS (full length approximately 1600bp). Based on the relative intensities of the bands, approximately half of the alleles in transfected LCLs had major deletions in the RBS. This resulted in a significant reduction in BFL-1 mRNA levels (**Figure 2E**). Transfecting either sgRNA individually led to limited editing, as determined by Sanger sequencing, but no significant difference in BFL-1 expression (**Figure 2 – Figure Supplement 1**). Thus, despite low H3K27ac levels, the RBS is an important NFκB-regulated node for BFL-1 expression.

### BFL-1 promotes resistance against mitochondria-dependent extrinsic apoptosis

NFκB signaling, which is activated downstream of LMP1, maintains homeostasis in EBV-immortalized LCLs by promoting the expression of pro- and anti-apoptotic proteins. For example, NFκB induces the expression of anti-apoptotic proteins BFL-1 and BCL-XL, which oppose pro-apoptotic sensitizers and effectors, while also upregulating the expression of proteins involved in extrinsic apoptosis, such as Fas and TRAIL receptors and their cognate antigens (Le Clorennec et al., 2008). Upon ligand binding, Fas/TRAIL receptors recruit and oligomerize FADD proteins into a death-inducing signaling complex (DISC) that activates caspase 8, the initiator caspase in extrinsic apoptosis (**Figure 3A**). Caspase 8 can directly activate executioner caspases 3/7 to induce mitochondria-independent apoptosis and can cleave the anti-apoptotic protein Bid into truncated Bid (tBid), a BH3-only peptide that induces intrinsic, mitochondria-dependent apoptosis. To prevent aberrant extrinsic apoptosis activation, NFκB upregulates the expression of c-FLIP, which prevents DISC formation and was shown to be critical for protecting against TNFα mediated extrinsic apoptosis in LCLs (Ma et al., 2017). This suggests that at steady-state, LMP1 expression in LCLs leads to constitutive activation of extrinsic apoptosis that is sufficiently inhibited by upregulated levels of anti-apoptotic factors.

**Figure 3.**
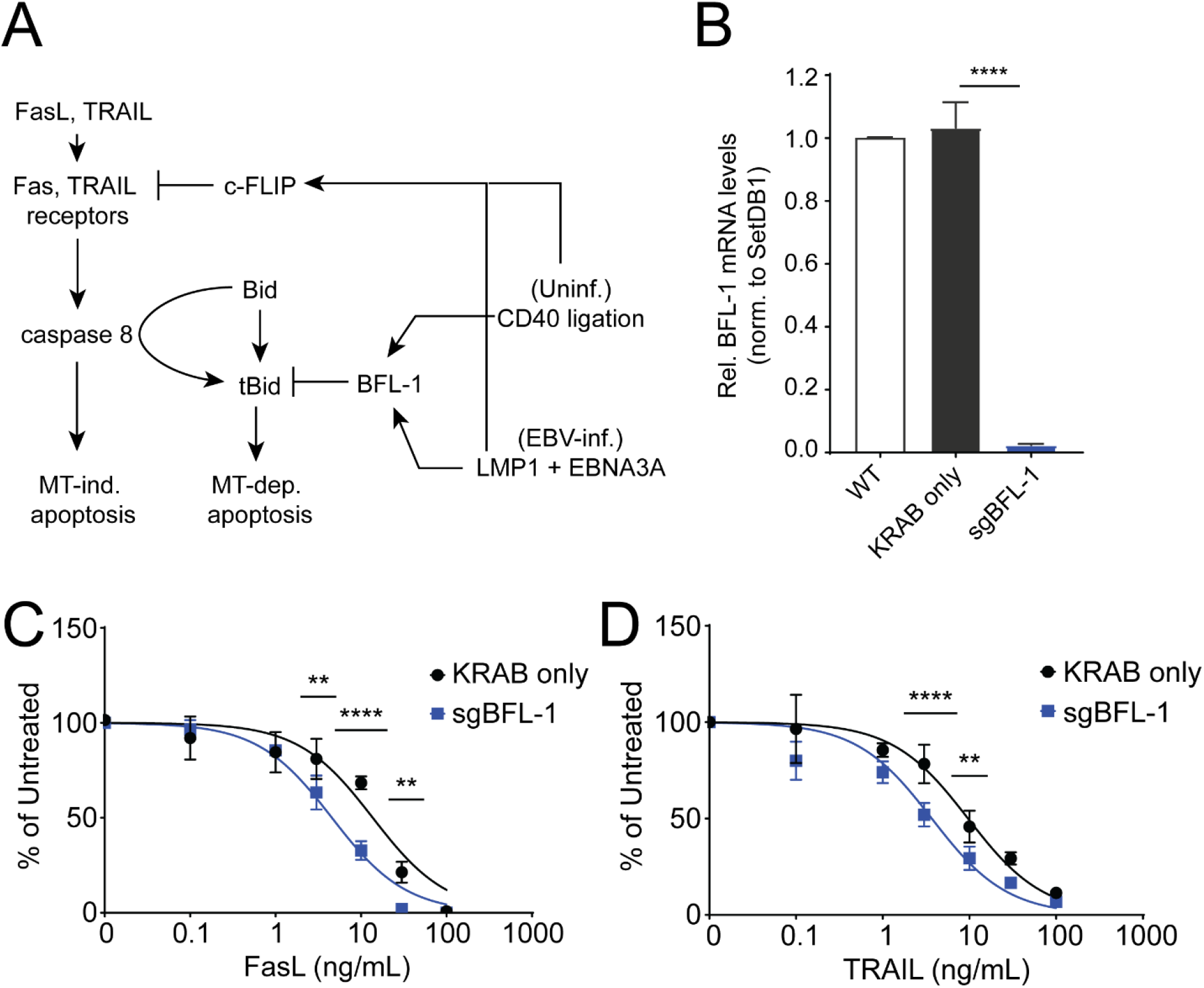
BFL-1 protects against extrinsic apoptosis. **A)** Schematic of FasL/TRAIL-mediated apoptosis of B cells in the germinal center light zone. NFκB signaling in the form of CD40 ligation in uninfected B cells or LMP1 expression in EBV-infected B cells induces expression of c-FLIP, which inhibits FasL/TRAIL-induced extrinsic apoptosis. Downstream, BFL-1 upregulation protects against mitochondrial-dependent apoptosis induced by tBid. **B)** qPCR of BFL-1 mRNA levels in WT LCLs and LCLs stably expressing dCas9-KRAB (only) and dCas9-KRAB targeted to the BFL-1 TSS (sgBFL-1). BFL-1 mRNA levels are normalized to SetDB1 and WT LCL. Mean and SEM from four experiments are reported. Significance determined by Two-way ANOVA, multiple comparisons to sgBFL-1. **** p < 0.0001. **C)** LCLs stably expressing dCas9-KRAB only (“KRAB only,” negative control) and dCas9-KRAB targeting the BFL-1 TSS (“sgBFL-1,” BFL-1 knockdown) were treated with TRAIL or **D)** FasL for three days and then assayed for cell counts relative to untreated control. Mean and SEM from two replicates are reported. Significance determined by Two-way ANOVA, comparing row cell means. ** p < 0.01, *** p < 0.001, **** p< 0.0001.

We hypothesized that BFL-1 upregulation in LCLs protects against mitochondria-dependent extrinsic apoptosis. We generated LCLs stably expressing dCas9-KRAB constructs that targeted the BFL-1 TSS, which significantly ablated BFL-1 mRNA levels (**Figure 3B**). BFL-1 knockdown LCLs were significantly more sensitive to increasing doses of FasL and TRAIL compared to negative control LCLs that expressed non-targeting dCas9-KRAB (**Figure 3C-D**). Therefore, these experiments performed on LCLs support a role for BFL-1 in protecting against mitochondrial-dependent extrinsic apoptosis.

### Epstein-Barr virus infection of primary human B cells *in vitro* phenocopies the germinal center reaction

Previously, we found that EBV-infected B cells and uninfected, maturing B cells undergo dynamic, temporal regulation of apoptosis (Dai and Luftig, 2018; Price et al., 2017). In particular, EBV-infected B cells and germinal center (GC) B cells upregulate strong dependence upon MCL-1 for survival, which led us to revisit the Germinal Center Model of *in vivo* EBV infection. Originally posited by David Thorley-Lawson, the GC Model posits that EBV latent infection is established in maturing B cells undergoing the GC reaction. The primary difference between *in vivo* and *in vitro* EBV infection is that EBV infection of B cells *in vitro* generates immortalized lymphoblastoid cell lines, or LCLs, that proliferate indefinitely in tissue culture (**Figure 4A**). However, EBV infection *in vivo* is primarily established in quiescent memory B cells. Primary EBV infection takes place in the oral cavity, where viral particles transmitted through the saliva first infect oral epithelial cells (**Figure 4B**). The infected epithelial cell amplifies the viral load thereby increasing the likelihood of infecting naïve B cells present in the oral mucosa. EBV activates and stimulates infected B cells to proliferate through the expression of viral proteins, which include six EBV nuclear antigens (EBNAs) and two latent membrane proteins (LMPs). The specific patterns of viral gene expression are categorized as “latency” types and are temporally regulated to induce proliferation, survival, immune evasion, and re-infection (reviewed in (Price and Luftig, 2015)).

**Figure 4.**
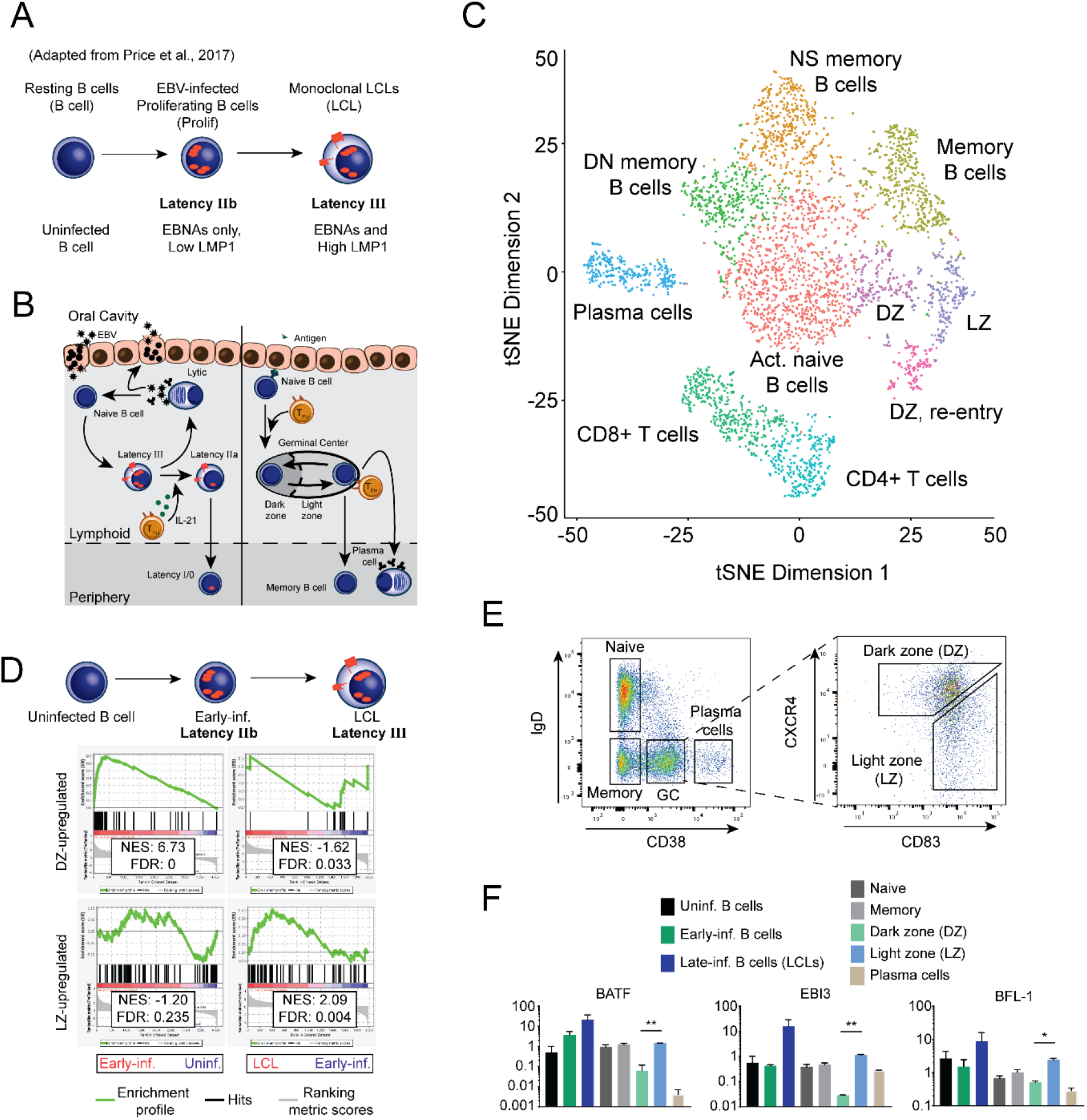
High BFL-1 expression typifies GC LZ B cells and EBV-immortalized LCLs. **A)** Schematic of EBV infection of human B cells *in vitro*. **B)** Schematic comparing EBV infection *in vivo* with B cell maturation in the oral cavity. Left – upon saliva transmission, EBV viral particles first infect an epithelial cell to amplify the initial viral load and increase the likelihood of infecting a naïve B cell in the lymphoid tissue. EBV infection activates the infected naïve B cell and stimulates it to proliferate through the expression of viral and host genes. First, EBV-infected B cells express the Latency IIb gene expression program, in which all EBV nuclear antigens (EBNAs) are expressed. Then, the EBNAs activate the expression of the latent membrane proteins (LMP), LMP1 and LMP2A, in Latency III. IL-21 secretion by T follicular helper cells (T_FH_) in the lymphoid follicle induces the transition to Latency IIa by silencing the expression of the EBNAs (Kis et al., 2010). Eventually, EBV-infected B cells attenuate viral gene expression in Latency 0, but periodically express EBNA1 in Latency I to maintain the viral genome in latently infected B cells. Plasma cell differentiation of EBV-infected B cells leads to lytic reactivation and production of infectious virion particles. Right – Affinity maturation of B cell antigen receptors is initiated when antigen encounter activates and stimulates naïve B cells to proliferate and form the GC reaction. In the GC dark zone (DZ), hyperproliferating B cells undergo class-switch recombination and somatic hypermutation and transit to the GC light zone (LZ) where they compete for antigen signaling and CD40 ligation from cognate T_FH_. Surviving B cells exit as plasma cells or as memory B cells. **C)** tSNE plot of scRNAseq of human tonsillar lymphocytes. Data represent two biological replicates that have been analyzed with canonical correlation analysis (CCA). NS – non-switched, DN – double negative. **D)** GSEA comparing differential gene expression between GC B cells and EBV-infected B cells *in vitro*. **E)** Flow cytometry plots of sorting strategy of CD19+ B cell subsets from tonsillar lymphocytes. **F)** qPCR comparing gene expression of BATF, EBI3, and BFL-1 in *in* vitro EBV-infected B cells and sorted tonsillar B cells. Data are representative of two experiments. Mean and SEM are plotted from two experiments and normalized to SetDB1 and uninfected B cells. Significance was determined by unpaired t-test between DZ and LZ B cells. * p < 0.05, ** p < 0.01.

From gene expression studies performed on tissue samples obtained from EBV-positive lymphomas and infectious mononucleosis, it was observed that EBV latency programs mimic gene expression in antigen-activated, maturing B cells undergoing the germinal center (GC) reaction (**Figure 4B**). The GC is spatially and functionally separated into two zones. In the dark zone (DZ), antigen-activated naïve B cells undergo rapid hyperproliferation, class switch recombination, and somatic hypermutation of antigen receptors (Victora et al., 2012). In the light zone (LZ), B cells compete for survival signals in the form of CD40 and BCR ligation from cognate T follicular helper cells. B cells with antigen receptors with subpar affinity are outcompeted and succumb to apoptosis (Victora et al., 2012). Surviving B cells exit as long-lived plasma cells and memory B cells or re-enter the DZ to undergo further affinity maturation (Dominguez-Sola et al., 2012; Dufaud et al., 2017).

To broadly characterize gene expression in maturing B cells, we performed single cell RNA-seq (scRNAseq) on cells isolated from human tonsillar lymphoid tissue (**Figure 4C**). T- and B-cell populations were identified based on relative levels of transcription factors and surface markers (**Figure 4 – Figure Supplement 1**). When plotted on a t-distributed stochastic neighborhood embedding (tSNE) plot, CD4+ and CD8+ T cells clustered together and away from CD19+ B cells (**Figure 4C, Figure 4 – Figure Supplement 1A-B)**. The plasma cell cluster, which expressed high levels of XBP1 and PRDM1/BLIMP1, was also distinctly separate from other B-cell populations, which clustered closely together and were distinguishable based on transcript levels of surface markers (**Figure 4 – Figure Supplement 1C**). We were able to identify GC B cell subsets based on relative expression of B-cell maturation factors and markers of proliferation and induction of the DNA damage response (**Figure 4 – Figure Supplement 1D-E**). Interestingly, our single cell gene expression analysis revealed a GC LZ cluster re-entering the GC DZ (**Figure 4 – Figure Supplement 1E**) (Dufaud et al., 2017). To confirm the accuracy of our identification of GC B cells, we performed a gene set enrichment analysis (GSEA) on GC DZ and LZ cluster markers with gene expression profiles generated from microarrays performed on sorted DZ and LZ populations (**Figure 4 – Figure Supplement 1F)**(Victora et al., 2012). This analysis, confirmed that GC B-cell clusters in our scRNAseq analysis were appropriately identified.

In our prior studies characterizing the early events after EBV infection, we identified several parallels between EBV infection and GC B cells. Similar to GC DZ B cells, early-infected B cells undergo rapid hyperproliferation, activation of the DNA damage response, and upregulation of anti-apoptotic dependency upon MCL-1 (Price et al., 2017). Combined with delayed upregulation of LMP1/NFκB signaling in LCLs, we surmised the early- and late-phases of *in vitro* EBV-infected B cells would be characterized by similar dynamic gene expression patterns in GC B cells. To determine a core set of genes that share similar dynamics of expression, we performed GSEA on differential gene expression between GC B cells and EBV-infected B cells. Using a pre-ranked list of DZ- or LZ-induced genes, we compared published microarray data that profiled gene expression changes in early- and late-infected B cells (Price et al., 2012). This analysis revealed that genes upregulated in early-infected B cells are enriched for targets induced in the DZ (**Figure 4D**). Conversely, genes upregulated in LCLs are preferentially expressed in the LZ. This analysis confirms that proliferation is a key defining trait for early-infected B cells and cycling GC DZ B cells, and LMP1/CD40 (NFκB) signaling are hallmarks for LCLs and GC LZ B cells.

Among the genes that made up the core enrichment of the GSEA, EBI3, BATF, and BFL-1 (BCL2A1), were strongly upregulated in GC LZ B cells and LCLs (**Figure 4 – Figure Supplement 1G**). To confirm this finding, qPCR was performed on sorted tonsillar B cells (**Figure 4E**) and EBV-infected B cells. LZ B cells were significantly more upregulated in EBI3, BATF, and BFL-1 mRNA levels compared to DZ B cells, as were LCLs compared to early-infected B cells (**Figure 4F)**. Using scRNA-seq, we were able to confirm that BFL-1 is robustly and uniquely upregulated in GC LZ B cells. In LCLs, we found that BFL-1 expression requires chromatin looping and epigenetic regulation by the viral nuclear protein EBNA3A (Price et al., 2017). We therefore sought to characterize the non-viral mechanisms underlying BFL-1 in GC LZ B cells.

### Chromatin in GC B cells is significantly more accessible and has increased levels of histone marks that characterize regions of active transcription and enhancer activity

To characterize the mechanisms by which BFL-1 becomes upregulated in uninfected GC LZ B cells, we first used ATAC-seq to identify regions of open chromatin (**Figure 5A**). Overall, we found that chromatin is significantly more accessible in GC subsets compared to naïve B cells, which confirms previous studies that show that B-cell activation induces significant chromatin opening to expose biologically active regions (**Figure 5 – Figure Supplement 1A**) (Bunting et al., 2016; Kieffer-Kwon et al., 2017). ATAC-seq also confirmed that naïve and memory B cells share highly similar chromatin landscapes (**Figure 5 – Figure Supplement 1B-C**). At *BCL2A1*, however, chromatin accessibility was not significantly different among B cell subsets; instead, upstream regions (demarcated “I” and “III”) were found to be significantly more open in GC B cells compared to naïve B cells (**Figure 5A**). However, both GC subsets shared similar chromatin openness despite different levels of BFL-1 expression, suggesting that additional mechanisms regulate LZ-specific induction of BFL-1 mRNA.

**Figure 5.**
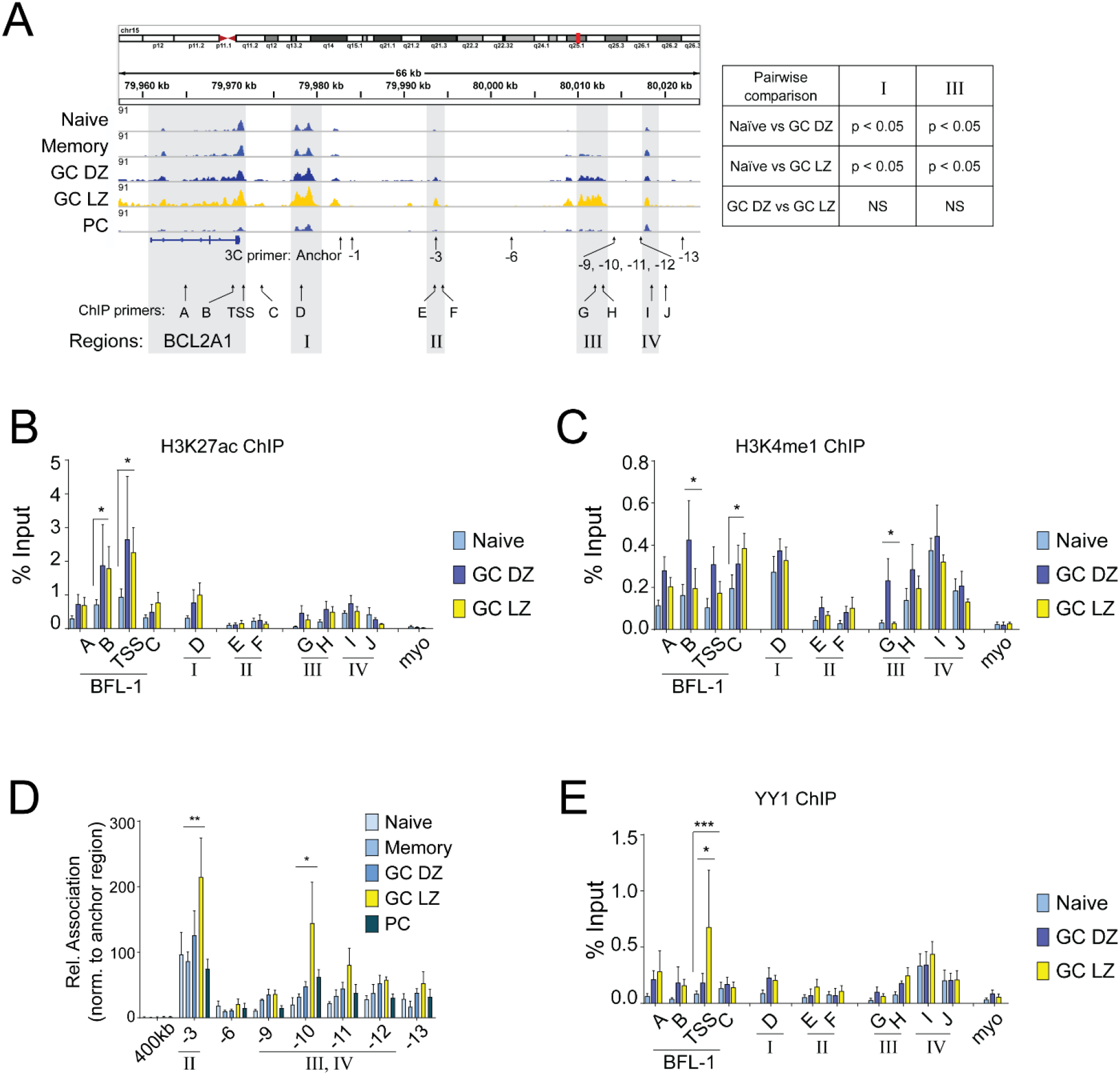
Increased association of open chromatin regions to the BFL-1 TSS in EBV-infected LCLs and GC LZ B cells. **A)** Representative ATAC-seq data from donor-matched CD19+ B cells sorted for naïve, memory, germinal center dark zone (GC DZ), light zone (GC LZ), and plasma cells (PC). Data shown are donor-matched subsets and are representative of three biological donors. Track heights are group auto-scaled and displayed in IGV using the hg38 assembly. Primers used for 3C-qPCR and ChIP-qPCR are listed below. Pairwise differential analysis on regions I and III performed on donor-matched replicates of ATAC-seq data from naïve, GC DZ, and GC LZ B cells. **B)** H3K27ac and **C)** H3K4me1 ChIP-qPCR performed on naïve, GC DZ, and GC LZ B cell subsets. Enrichment calculated relative to input. Myoglobin (myo) was included as a negative control. Mean and SEM from three experiments are plotted. Significance determined by Two-way ANOVA, multiple comparisons to the GC LZ population. * p < 0.05, ** p < 0.01, *** p < 0.001. **D)** 3C-qPCR performed on sorted naïve, memory, GC DZ, GC LZ, and plasma cells from tonsillar CD19+ B cells. HindIII-produced fragments of chromatin were interrogated for interactions with the anchor primer near the BFL-1 TSS. Interaction frequencies were normalized to those of the nearest neighbor HindIII fragment, −1. Mean and SEM from three experiments are plotted. Significance determined by Two-way ANOVA with Holm-Sidak’s multiple comparisons test. **** p < 0.0001, ** p < 0.001. The HindIII fragment located 400kb from the BFL-1 TSS served as a negative control. **E)** YY1 ChIP-qPCR performed on naïve, GC DZ, and GC LZ B cell subsets with enrichment normalized to input. Mean and SEM from three experiments are plotted. Significance determined by Two-way ANOVA, multiple comparisons to the GC LZ population. * p < 0.05, ** p < 0.01, *** p < 0.001.

We next sought to determine if chromatin regions were differentially activated at the BFL-1 locus among B-cell subsets. H3K27ac levels were elevated in GC DZ and LZ subsets and more so at the BFL-1 gene body than at the enhancers (**Figure 5B**). B-cell subsets shared similar levels of H3K4me1 levels but were preferentially higher in GC DZ B cells near the BFL-1 TSS, suggesting poised transcription (Bae and Lesch, 2020). H3K4me1 is also moderately enriched at Enh 2, which suggests poised enhancer activation (**Figure 5C**). These data indicate that increased chromatin accessibility in GC B cell subsets is accompanied by increased levels of enhancer activity.

To characterize the chromatin interactions at the BFL-1 locus in GC LZ B cells, we performed chromatin conformation capture (3C) analysis on sorted B-cell subsets. 3C-qPCR revealed cell type-specific chromatin structures upstream of the BFL-1 gene in which interactions between region “I” and regions “II” and “III, IV” were significantly higher in GC LZ B cells than in any other B cell subset (**Figure 5D**). Therefore, while GC DZ and LZ B cells share similar levels of chromatin accessibility, the formation of a specific three-dimensional chromatin architecture facilitates BFL-1 upregulation in GC LZ B cells. Interestingly, this architecture closely resembles the looping found in EBV-immortalized LCLs (**Figure 1D**). Since we had found that EBNA3A-null LCLs had reduced levels of chromatin-bound YY1, we also performed ChIP-qPCR in sorted B cell subsets for YY1, which plays a critical role in all stages of B cell development and regulates the germinal center transcription program (Green et al., 2011; Kleiman et al., 2016). Among queried naïve, GC DZ, and LZ populations, YY1 occupies the chromatin landscape at similar levels, but was uniquely and significantly higher at the BFL-1 TSS in GC LZ B cells (**Figure 5E**). This suggests that YY1 is uniquely recruited in GC LZ B cells to facilitate the looping of activated enhancers to the BFL-1 TSS. Overall, these data show that BFL-1 transcription in GC LZ B cells occurs through stage-specific priming, looping, and activation of enhancers during B-cell maturation.

## DISCUSSION

Here, we have shown parallels between BFL-1 upregulation in EBV-immortalized LCLs and in uninfected GC LZ B cells. BFL-1 is one of several shared targets that are upregulated in EBV-infected B cells and GC LZ B cells. Previously, it was shown in LCLs that LMP1, EBNA2, and EBNA3A were important for BFL-1 transcription. LMP1-induced NFκB signaling upregulates BFL-1 expression (D’Souza et al., 2004), EBNA2 activates BFL-1 transcription at the TSS (Pegman et al., 2006), and EBNA3A is required for looping distal enhancers to the BFL-1 TSS (Price et al., 2017). Loss of EBNA3A severely abrogates BFL-1 expression, which is still maintained at low levels due to EBNA2 and LMP1 activities. BFL-1 is therefore an important viral target in EBV infection, and its expression depends upon coordination among multiple viral proteins.

In GC B cells, chromatin accessibility at the BFL-1 locus and upstream regions becomes significantly more open and share similar levels of H3K27ac. Increased chromatin interactions and YY1 binding at the BFL-1 TSS in GC LZ B cells promote BFL-1 transcription and a chromatin architecture that resembles that in WT LCLs. A direct relationship between EBNA3A and YY1 has not been shown previously, although both are important in mediating ESE activity. However, as a polycomb group protein, YY1 may be recruited to EBNA3A-occupied sites to mediate chromatin looping and gene expression (Hickabottom et al., 2002; Skalska et al., 2010; Srinivasan and Atchison, 2004; Touitou et al., 2001). In LCLs, functional interrogation of genomic regions using CRISPR and CRISPRi shows that BFL-1 expression is controlled by active enhancers and interacting domains. The similarities in chromatin regulation between LCLs and GC LZ B cells suggest that BFL-1 is an important target for both B cell maturation and EBV infection.

The Germinal Center (GC) Model posits that B cells infected by EBV *in vivo* must transit through the germinal center reaction to gain access to the long-lived memory B cell compartment. Our data supports this model by showing that an *in vitro* EBV infection of B cells phenocopies several important facets of GC B cell biology. The GC reaction is functionally and spatially segregated into two zones, which are mimicked by the early- and late-phases of EBV infection. Both DZ B cells and early-infected B cells undergo rapid hyperproliferation and induction of the DNA damage response, while LZ B cells and LCLs grown out from the late-phase of EBV infection are characterized by CD40/BCR signaling. Transcriptionally, early-infected B cells are enriched for DZ-upregulated genes and LCLs are enriched for LZ-upregulated genes, such as critical B cell maturation factors like IRF4 and BATF that are essential for LCL survival (Ma et al., 2017). These observations indicate that many aspects of the GC reaction, such as dynamic regulation of transcription and chromatin regulation, are intrinsic to EBV infection.

We therefore present an updated version of the GC Model, based on this work and others from our lab (**Figure 6**). In the GC DZ, EBV-infected B cells express the Latency IIb program, which drives proliferation. Upon entering the GC LZ, EBV-infected B cells express LMP1 and LMP2A in the Latency III program to promote survival. Originally, the GC model suggested that EBV-infected B cells expressed Latency III and IIa in the GC, in which LMP1 and LMP2A promote proliferation and survival of the infected reservoir. However, LMP1 expression could be deleterious in that it induces potent cytotoxic T cell responses (Choi et al., 2018). In addition, EBV-positive Burkitt lymphomas, which express high c-Myc but no LMP1, are relatively non-immunogenic (Staege et al., 2002). Therefore, the Latency IIb program would permit rapid expansion of infected B cells with relatively low immunogenicity. This new GC model shows that the temporal regulation of viral gene expression observed in B cells infected *in vitro* is in accord with GC B cell dynamics and biphasic NFκB activity.

**Figure 6.**
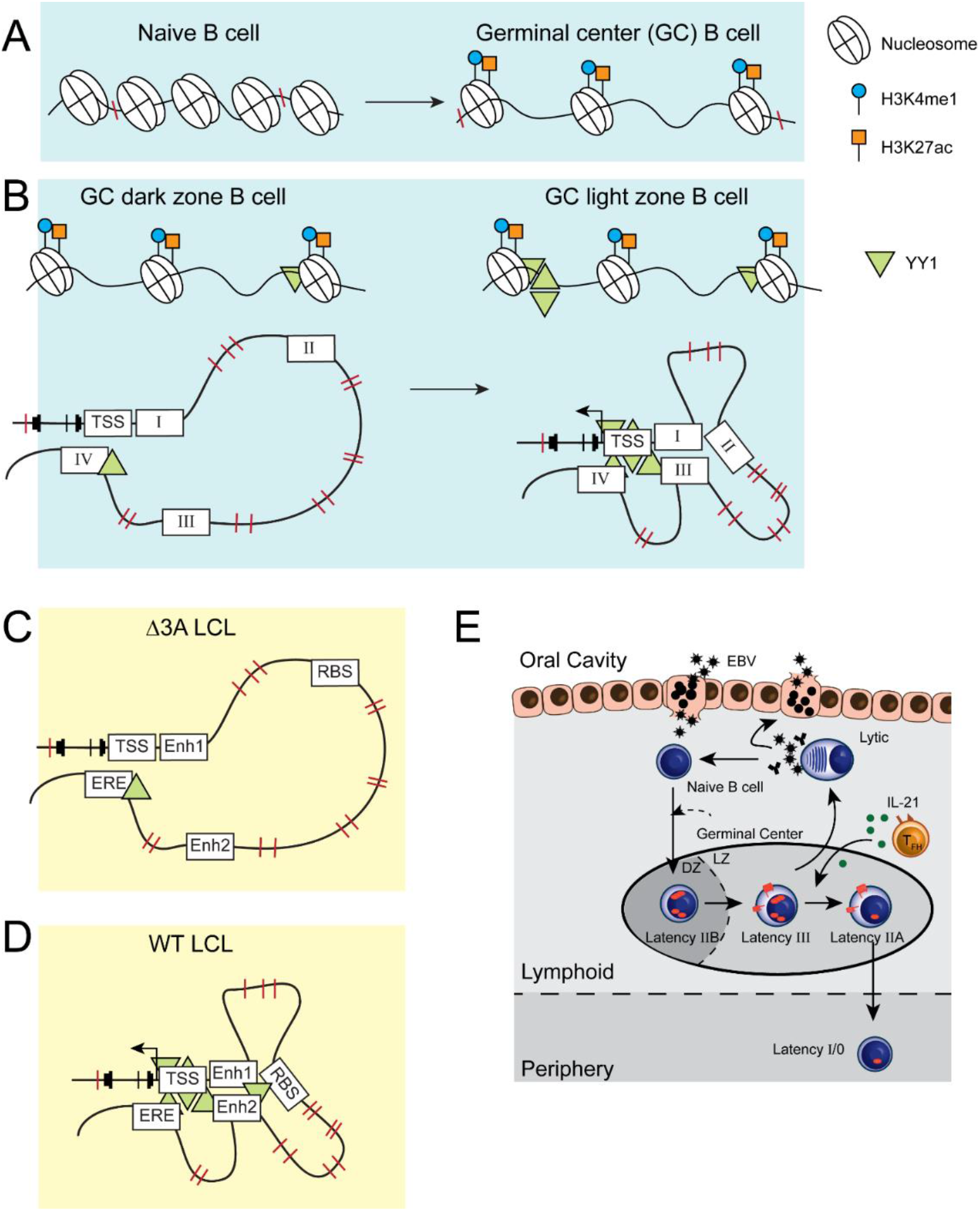
Illustrated schematic of stage-specific chromatin accessibility, activation, and looping in maturing B cells and EBV-infected B cells. **A)** The transition from naïve to GC B cell is characterized by increased chromatin accessibility at *BCL2A1* and at enhancer regions. **B)** At *BCL2A1*, increased YY1 deposition at the chromatin in GC LZ B cells facilitates the looping of activated and accessible enhancers in a specific chromatin architecture. This allows for LZ-specific upregulation of BFL-1 transcription. **C)** EBNA3A-null LCLs have lower levels of chromatin-bound YY1 and lack the chromatin looping observed in **D)** wildtype (WT) LCLs. Importantly, the chromatin architecture in WT LCLs strongly resembles that in GC LZ B cells. **E)** A new, updated version of the GC Model of EBV infection incorporates the gene expression similarities, anti-apoptotic dependencies, and chromatin regulation observed in *in vitro* models of EBV infection to those observed in maturing B cells.

However, there are important differences between GC LZ B cells and EBV-immortalized LCLs. For example, at the BFL-1 locus, H3K27ac was preferentially enriched at upstream enhancers in LCLs whereas H3K27ac levels were higher at the BFL-1 gene than at enhancers in GC LZ B cells. This most likely reflects the fact that these viral-regulated enhancers control multiple targets. RNA pol II ChIA-PET shows that Enh 2 and the ERE are strongly linked to both BFL-1 and an upstream gene, *ZFAND6*, which is an LCL-essential gene (Ma et al., 2017). Also known as AWP1, ZFAND6 modulates NFκB activity (Chang et al., 2011; Fenner et al., 2009), but a role for ZFAND6 in GC B cells has not been identified, despite the importance of NFκB signaling in GC LZ B cells. It is possible that ZFAND6 may be more important for regulating sustained NFκB signaling in LCLs rather than in transient settings such as the GC LZ.

Other important discrepancies involve BCL6 and c-Myc expression, both of which are required for the GC reaction. B cells infected with EBV *in vitro* strongly downregulate BCL6 expression (Pei et al., 2017; Price et al., 2012), suggesting perhaps that EBV infection is incompatible with the GC. However, EBV-infected B cells found *in vivo* can express BCL6, indicating that BCL6 expression is dependent upon environmental cues. While both early-infected B cells and LCLs express c-Myc (Nikitin et al., 2010), c-Myc is rarely expressed in GC B cells but is nonetheless required for GC formation and plays a critical role in mediating chromatin changes in activated B cells (Dominguez-Sola et al., 2012; Kieffer-Kwon et al., 2017). Levels of c-Myc are upregulated among GC LZ B cells that re-enter the DZ to undergo further affinity maturation and their duration in the DZ depends upon the initial levels of c-Myc in the returning GC LZ B cell (Finkin et al., 2019; Heinzel et al., 2017). For an EBV-infected B cell, c-Myc is important for maintaining the latently-infected state (Guo et al., 2020), but high c-Myc levels could lead to excessive retention in the GC and an increased risk of being detected and eliminated by T cells. EBV-infected B cells may overcome this challenge by restricting viral expression to Latency IIa, which occurs in response to IL-21 secretion by T_FH_ (Kis et al., 2010). IL-21 silences EBNA2, the primary activator of c-Myc expression, and the EBNA3s, which are highly immunogenic. Thus, EBV-infected B cells are equipped to respond to appropriate cues and to overcome the barriers of the GC reaction.

The study of EBV infection *in vivo* has been challenging because primary EBV infection is often asymptomatic, and therefore difficult to observe, and the frequency of infected B cells in asymptomatic cases is extremely low. As a result, much of what we know about *in vivo* infection has been informed by painstaking single-cell PCR experiments for viral transcripts and inferred from studies of EBV-associated diseases and tumors. The advent of scRNA-seq and improved chromatin technologies promises a better understanding of how EBV establishes latent infection in healthy individuals and how this becomes dysregulated in EBV-associated malignancies.

## MATERIALS AND METHODS

### Cell Culture

LCLs used in this study were also used in Price et al., 2017. In short, peripheral blood mononuclear cells (PBMCs) were harvested from human donors through the Gulf Coast Regional Blood Center (Houston, TX). CD19+ B cells were isolated from PBMCs using the BD iMag Negative Isolation Kit (BD, 558007) and subsequently infected with the wildtype (WT) B95-8 Epstein-Barr virus strain as previously described (Johannsen et al., 2004). The recombinant EBNA3A-null virus that was used in this study and in (Price et al., 2017) was generated in (Bazot et al., 2015). LCLs were maintained in RPMI supplemented with 10% FBS (Corning), 2mM L-Glutamine, 100U/mL penicillin, and 100μg/mL streptomycin (Invitrogen).

Tonsillar B cells were isolated from discarded, anonymized tonsillectomies from the Duke Biospecimen Repository and Processing Core (BRPC; Durham, NC). Tonsil tissue samples were manually disaggregated, filtered through a cell strainer, and isolated by layering over a cushion made from Histopaque-1077 (H8889; Sigma-Aldrich). Harvested lymphocytes were washed three times with FACS buffer (5% FBS in PBS) and stained for surface markers using: CD19-PE (363003, Biolegend, RRID: AB_2564125), IgD-FITC (348206, Biolegend, RRID: AB_10612567), CD38-APC (303510, Biolegend, RRID: AB_314362), CD83-BV421 (305324, Biolegend, RRID: AB_2561829), CXCR4-PeCy7 (560669, BD Biosciences, RRID: AB_1727435).

For flow cytometry analysis, stained cells were analyzed on a BD FACS Canto II and sorted on a MoFlo Astrios Cell Sorter at the Duke Cancer Institute Flow Cytometry Shared Resource (Durham, NC).

### Chromatin Immunoprecipitation (ChIP)

ChIP was performed as described previously (Bazot et al., 2015) using an Active Motif ChIP-IT PBMC assay kit (Cat. No. 53042). Cells were fixed for 8 min and sonicated for 60 min (30 sec on, 30 sec off) on high with a standard Diagenode Bioruptor Sonicator. Sonicated ChIP input was reverse-crosslinked and purified to run on a 1.5% agarose gel to ascertain appropriate fragment sizes. Antibodies used were: Rabbit anti-YY1 (Active Motif, 61779, RRID:AB_2793763), Rabbit anti-H3K27ac (Active Motif, 39135, RRID:AB_2614979), Rabbit anti-H3Kme1 (EMD Millipore, 07-436), Rabbit IgG isotype control (Thermo Fisher, 02-6102)

### Chromatin Conformation Capture (3C)

3C was performed as previously described (Hagege et al., 2007). In brief, chromatin was crosslinked with formaldehyde and isolated from LCLs, whereupon it was digested overnight with HindIII-HF (NEB, R3104M). Digested chromatin was then diluted and ligated overnight at 16°C with T4 DNA ligase (NEB, M0202M). Reversal of crosslinks, proteinase K digestion, and phenol-chloroform extraction yielded DNA circles that were assayed by TaqMan qPCR. Ct values were normalized to enrichment to a region closest to the anchor primer.

### ATAC-seq

Three biological replicates of sorted CD19+ B cells (naïve, memory, GC DZ, GC LZ, and plasma cells) were isolated and processed for ATAC–seq as described in (Buenrostro et al., 2015). Samples were run on 2 lanes of the Illumina 4000 HiSeq. ATAC-seq data was processed as previously described. Adaptor sequences were trimmed and mapped to hg38 using Bowtie2. Reads were then filtered for duplicates. The filtered reads for each sample were merged, and peak calling was performed by MACS2.

### dCas9-KRAB systems

A short-guide RNA targeting the BFL-1 transcriptional start site was cloned into the BsmbI-digested dCas9-KRAB-GFP (Addgene #71237, RRID: Addgene_71237) or lentiguide-puromycin (Addgene #52963, RRID: Addgene_52963) vector. Ligated vector was then transformed into Stbl2 and sequence confirmed with the hU6 primer. 1μg plasmid was then transfected with 1μg psPAX2 (Addgene #12260, RRID: Addgene_12260) and 100ng pMD2.G (Addgene #12259, RRID: Addgene_12259) in 293Ts with Mirus LTI reagent in antibiotic-free media and harvested 48 and 72 hours post transfection. Virus was then filtered and concentrated prior to transducing LCLs. LCLs transduced with dCas9-KRAB-GFP were then sorted for GFP+ events and grown out. LCLs transduced with lentiguide-puromycin were selected with 0.8 μg/mL puromycin for 4-7 days and then transduced with TRE-HAGE-dCas9-KRAB lentivirus, which was generated similarly. Transduced cells were then selected with 100μg/mL G418 in RPMI supplemented with 15% Tet-free FBS. To induce TRE-HAGE-dCas9-KRAB activity, cells were treated with 3μg/mL doxycycline for 48hrs and then harvested for mRNA. To show Dox-specific effects, treated cells were washed twice with Tet-free media and cultured for 48hrs before harvesting for mRNA.

### Cas9 RNP transfection

Cas9 RNP transfections were performed based on manufacturer’s instructions (Thermo Fisher Scientific, TrueCut Cas9 Protein v2). sgRNAs were synthesized by Synthego. In short, Cas9 protein and sgRNAs were mixed in a ratio of 1:3 and incubated in Belzer’s solution for 20min at room temperature. Cells were used at a final concentration of 10 million/mL. Microporations were performed with the Neon system with 10uL tips. Cells were recovered in RPMI supplemented with 15% FBS and no antibiotic. 24 hours after electroporation, cells were supplemented with 1% Penicillin-Streptomycin-Glutamine. mRNA and genomic DNA were harvested 5 days post transfection to assay for BFL-1 mRNA levels by qPCR and cleavage of the genomic DNA by PCR, which was performed with GoTaq master mix (Promega).

### Single cell RNA-seq

Two biological replicates of tonsillar lymphocytes were prepared for single cell RNA-seq using technology from 10X Genomics. Libraries were run on one lane of the Illumina 4000 HiSeq. Reads were processed, aligned, and normalized using Cell Ranger from 10X Genomics.

Processed scRNAseq were analyzed using the Seurat R package (version 2.2) (Butler et al., 2018) for graph-based clustering and visualizations. Processing of data was performed with the default parameters. Cells that passed quality control processing and expressed at least 200 genes and only genes that were expressed in at least 3 cells. Cells with greater than 5% mitochondrial genes were removed from analysis.

Both tonsillar lymphocyte replicates were combined and analyzed using Seurat’s Canonical Correlation Analysis (CCA) with RunCCA to identify conserved gene correlations. We used the top 2000 variable genes from each sample to calculate the correlation components (CCs) and determined that 11 CCs sufficiently represented the variability in the dataset. Both samples were then aligned with AlignSubspace using 11 CC dimensions. FindClusters was then used to apply SNN clustering to the combined cells using the 11 aligned CCs and resolution 0.5. Differentially expressed genes were determined prior to clustering, which was visualized with t-distributed stochastic neighbor embedding (tSNE) dimensionality reduction using RunTSNE (11 aligned CCs) and TSNEPlot. Seurat object and analysis were updated using the UpdateSeuratObject function.

### Quantitative PCR

mRNA was isolated from samples using the Promega SV 96 Total RNA Isolation System and reverse-transcribed into cDNA using the High-Capacity cDNA Reverse Transcription Kit from Thermo Fisher. qPCR was performed using the Power SYBR Green PCR Master Mix from Thermo Fisher. Relative mRNA values were calculated using the ΔΔCt method and normalizing to SetDB1 or ALAS1 housekeeping genes.

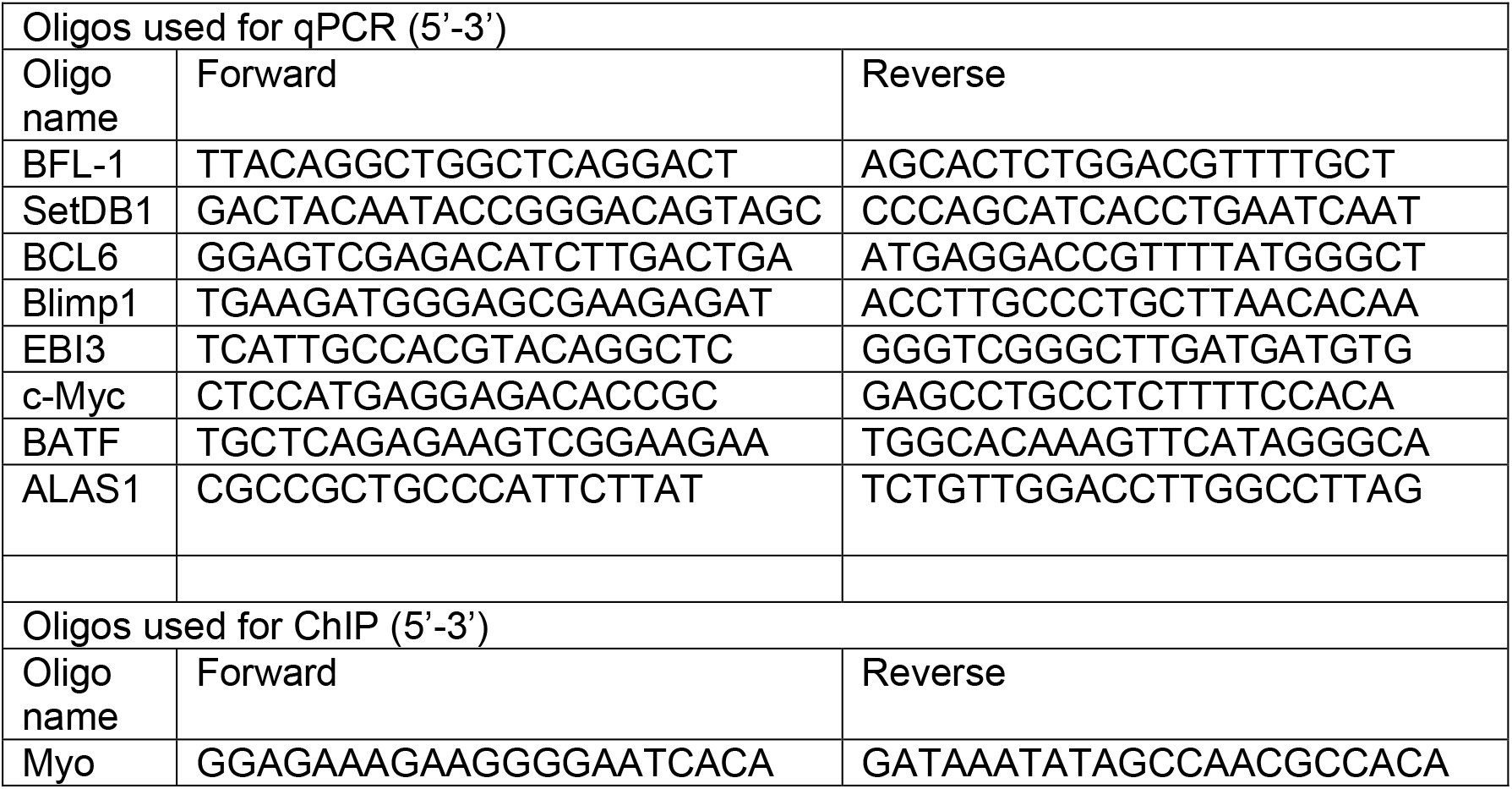
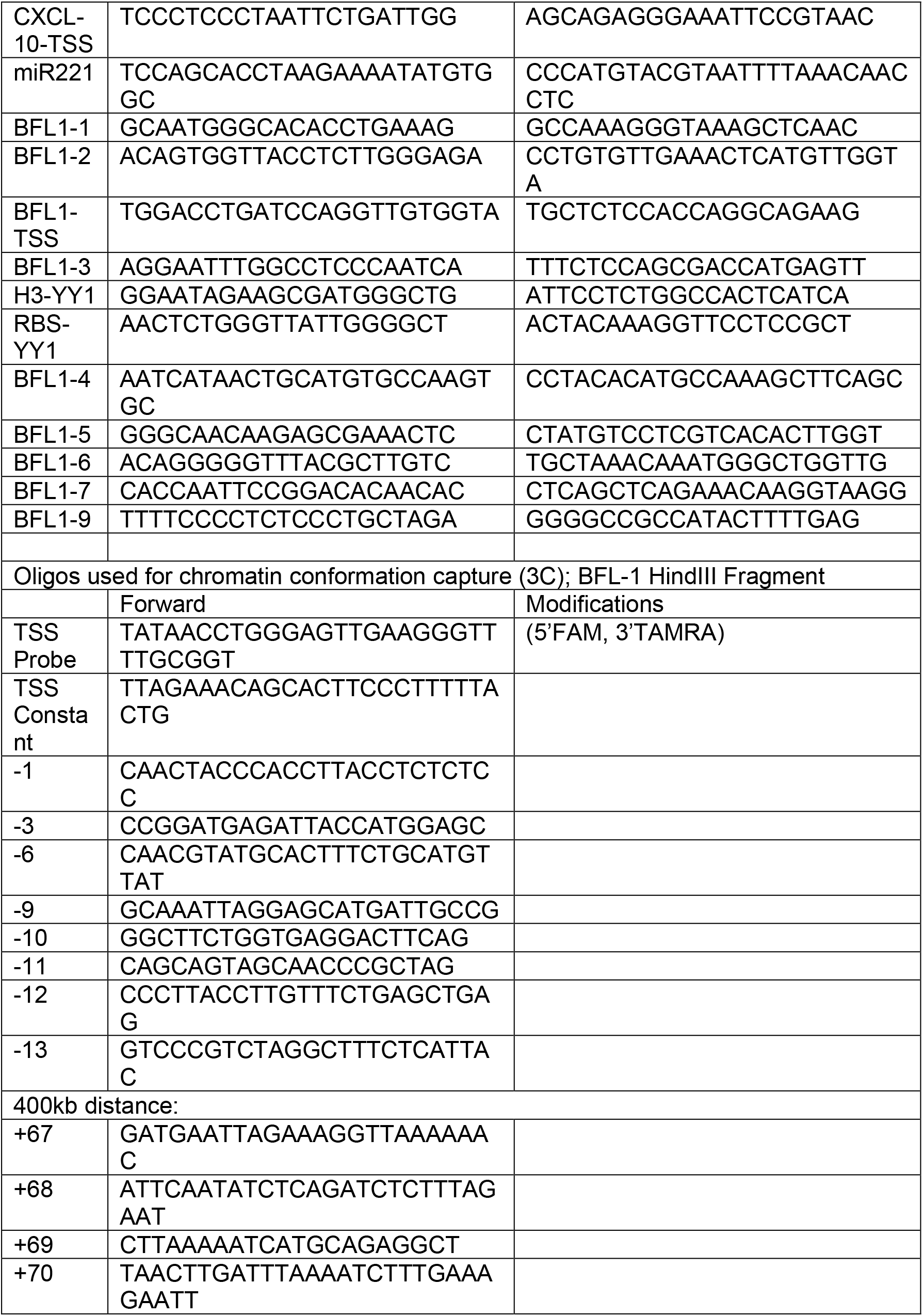
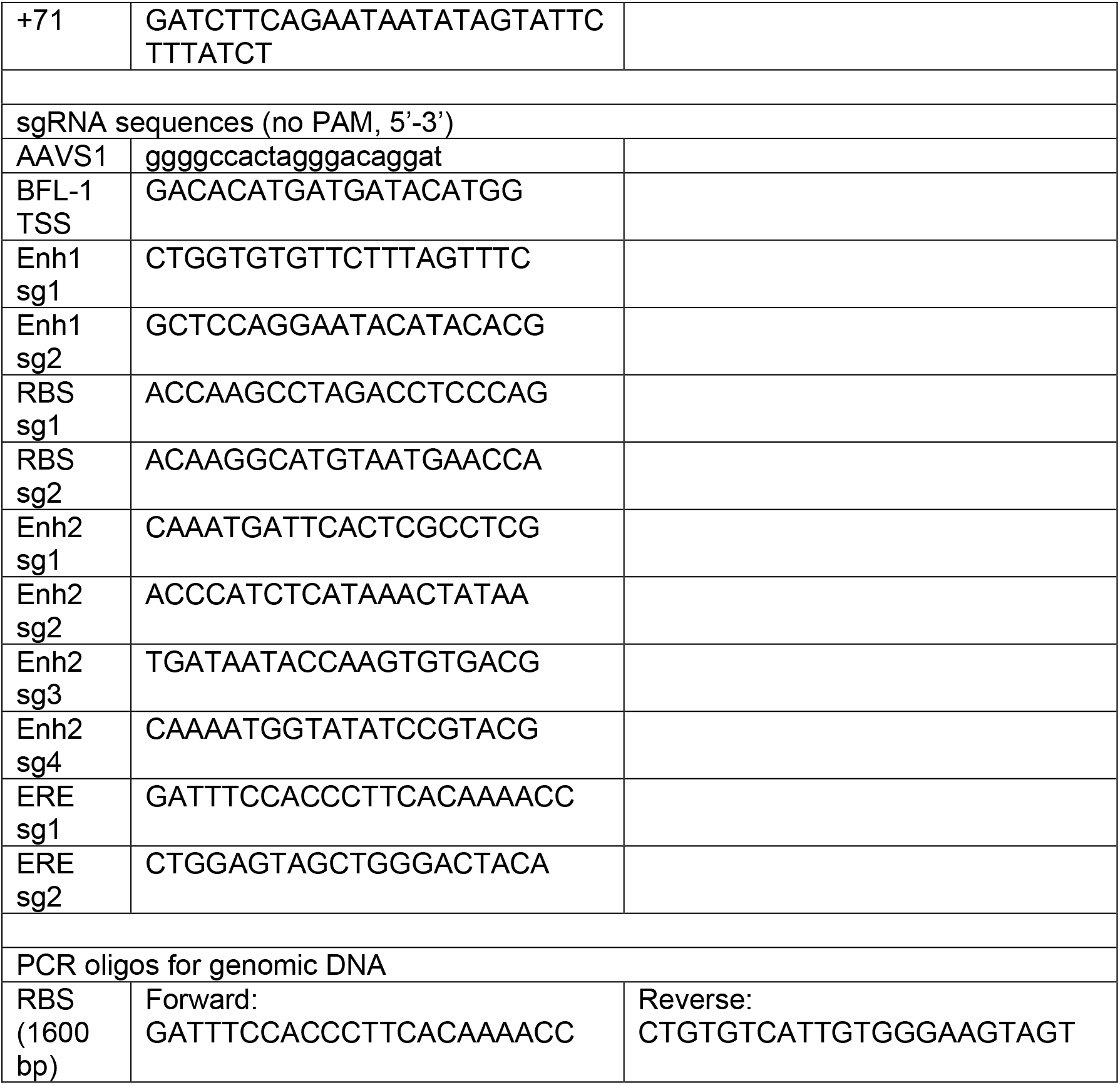

---

**Figure 2 - Figure Supplement 1.**
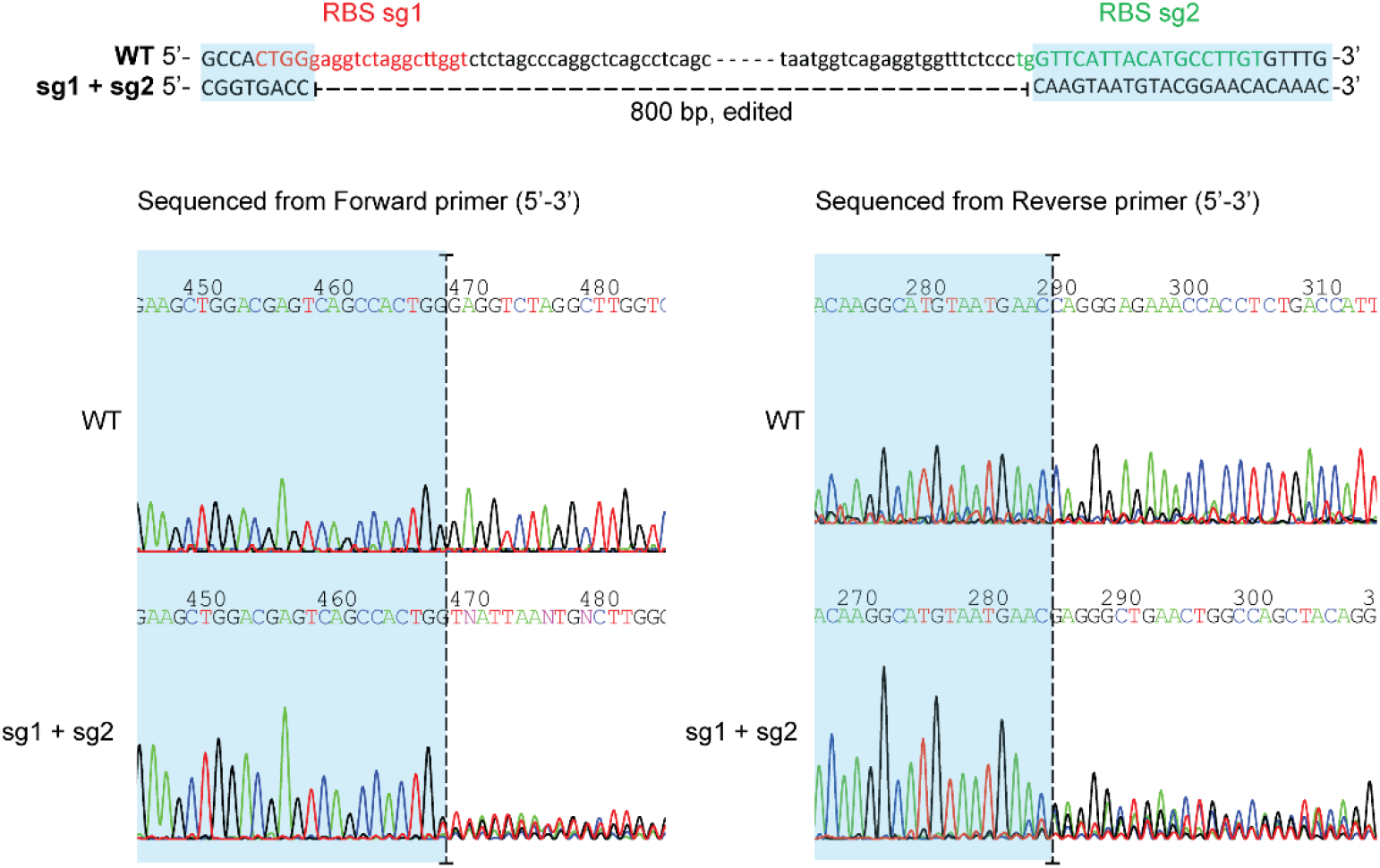
Bulk Sanger sequencing of edited RBS genomic region. Due to major sequence alterations and deletions, editing efficiency was not quantifiable by Synthego ICE (Inference of CRISPR Edits) analysis. Trace files shown below. Sequences highlighted in blue are aligned.

**Figure 4 – Figure Supplement 1.**
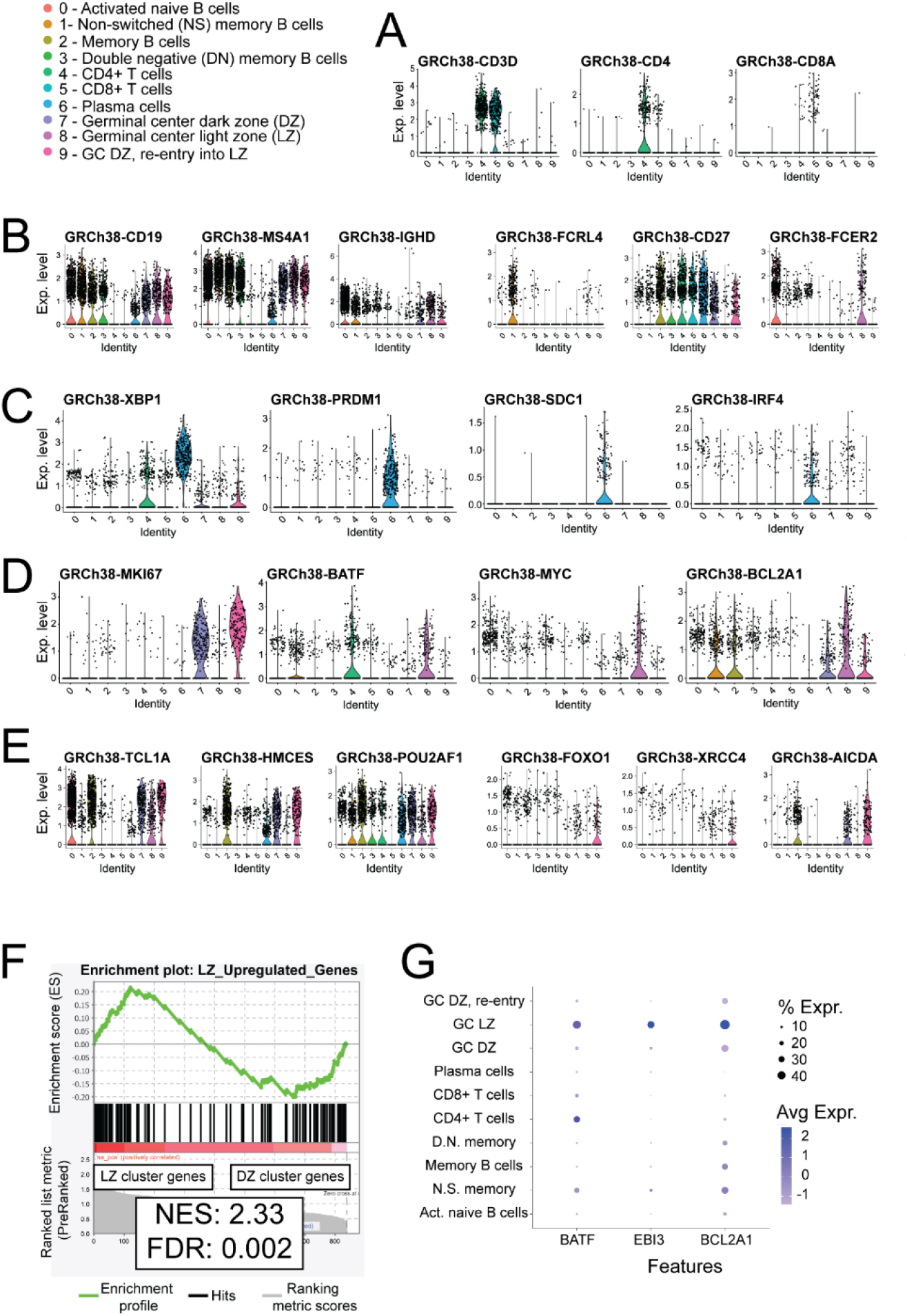
Single cell RNA-seq (scRNA-seq) identifies markers of T and B lymphocyte clusters in human tonsillar tissue. **A)** Violin plots generated for CD3+ (CD3D+) T cell clusters. Cluster 4 – CD4+ T cells, Cluster 5 – CD8+ T cells. GRCh38 – human genome assembly 38. **B)** Violin plots generated for naïve and memory B cell clusters. Cluster 0 – Activated naïve B cells, Cluster 1 – Non-switched (NS) memory B cells, Cluster 2 – Memory B cells, Cluster 3 Double negative (DN; IgD-low, CD27-low memory B cells). **C)** Violin plots generated for Cluster 6 – plasma cells. **D)** Violin plots generated for Cluster 8 – GC LZ B cells. **E)** Violin plots generated for Cluster 7 – GC DZ B cells, and Cluster 9 – GC DZ B cells returning from the LZ. Identities based on (Dufaud et al., 2017). **F)** GSEA comparing differentially expressed genes in GC LZ and DZ B cell clusters from scRNAseq and with previously published data in (Victora et al., 2012). Dot plot of scRNA-seq data of relative expression levels of LZ-upregulated markers BATF, EBI3, and BCL2A1 among lymphocyte clusters.

**Figure 5 - Figure Supplement 1.**
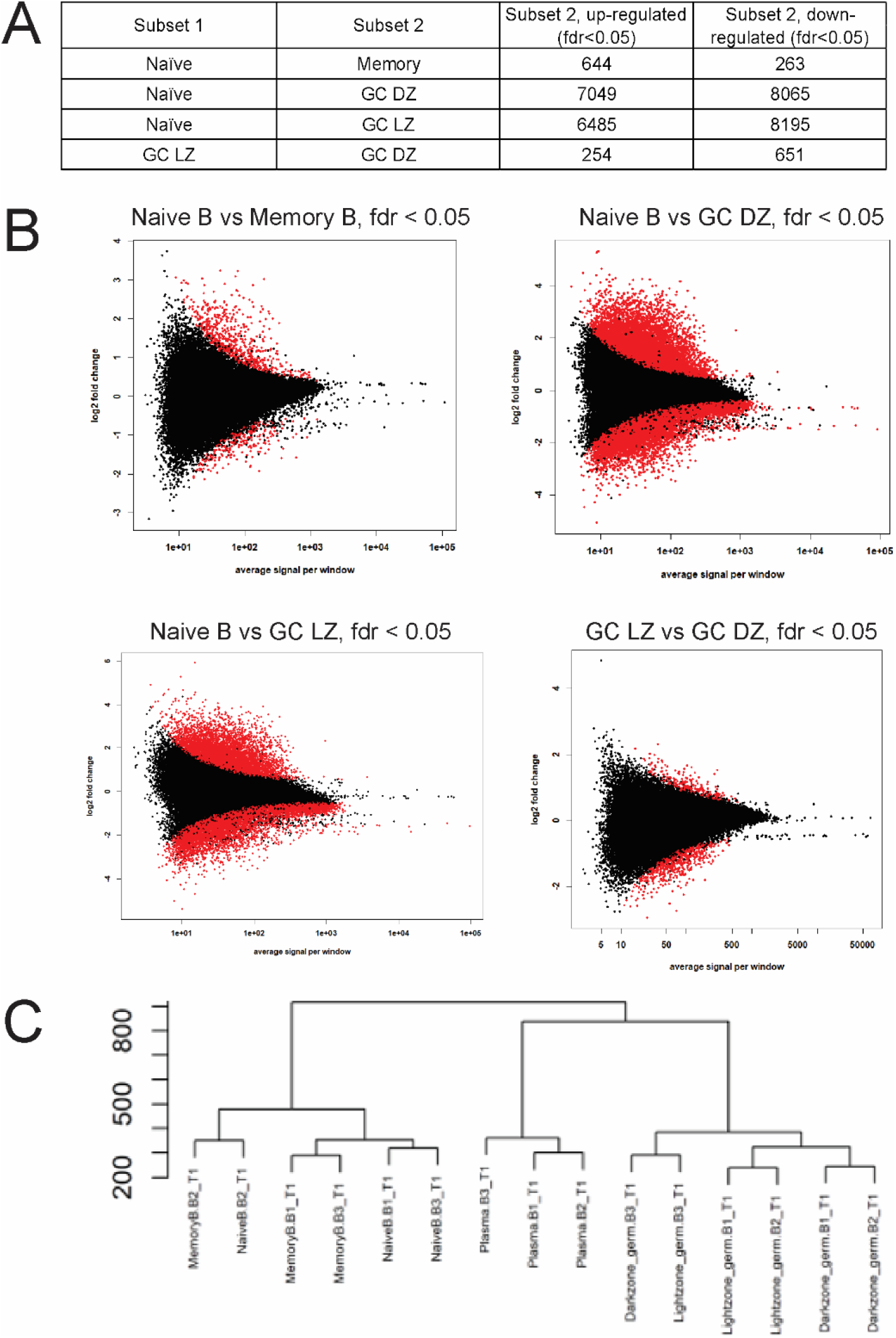
Differential analysis of open chromatin regions in B cell subsets. **A)** Pairwise t differential analyses on B cell subsets from three biological donors were processed by DESeq2 with cut off false discovery rate (fdr) < 0.05 (Love et al., 2014). **B)** MA plots (log2 fold change vs average signal per peak) shows changes in chromatin accessibility for all peaks. Red dots indicate **C)** Hierarchical clustering shows strong similarities between naïve and memory B cell subsets and within GC B cell subsets.

## ACKNOWLEDGMENTS

This work was supported by National Institute of Health (NIH) grants R01-CA140337 and R01-DE025994 (to M.A.L.), T32-CA009111 and F31-DE027875 (to J.D.). We would like to acknowledge the assistance of the Duke Molecular Physiology Institute Molecular Genomics Core for the generation of the scRNA-seq in this manuscript. The authors would also like to thank Dr. Michael Cook, Nancy Martin, Lynn Martinek and the Duke Cancer Institute Flow Cytometry Shared Resource for invaluable assistance with flow cytometry and cell sorting. Finally, the CRISPR work was greatly assisted by Dr. So Young Kim in the Duke Cancer Institute Functional Genomics Shared Resource.

## AUTHOR CONTRIBUTIONS

J.D. and M.A.L. conceived and designed the research. J.D. executed and oversaw all of the experiments in this manuscript except for the ATAC-seq. E.H. assisted with the ChIP-QPCR experiments. L.S. performed the ATAC-seq experiments and analysis. G.E.C. interpreted ATAC-seq experiments and edited the manuscript. J.D. and M.A.L. wrote and edited the manuscript.

